# Annotation and assessment of functional variants in regulatory regions using epigenomic data in farm animals

**DOI:** 10.1101/2024.02.06.578787

**Authors:** Ruixian Ma, Renzhuo Kuang, Jingcheng Zhang, Jiahao Sun, Yueyuan Xu, Xinbo Zhou, Zheyu Han, Mingyang Hu, Daoyuan Wang, Yuhua Fu, Yong Zhang, Xinyun Li, Mengjin Zhu, Shuhong Zhao, Tao Xiang, Mengwei Shi, Yunxia Zhao

## Abstract

**Background:** Understanding the functional impact of genetic variants is essential for advancing animal genomics and improving livestock breeding. Variants that disrupt transcription factor (TF) motifs provide a means to assess functional potential, but the lack of TF ChIP-seq data for farm animals presents a challenge.

**Results:** To address this, we curated nearly 900 epigenomic datasets from 10 farm animal species and annotated eight regulatory regions to assess how variants affect TF motifs. Over 127 million candidate functional variants were classified into five functional confidence categories across the species. Variants with high confidence were enriched in eQTLs and trait-associated SNPs, showing greater potential to affect gene expression and phenotypes. Incorporating these functional variants into genomic prediction models improved the accuracy of Estimated Breeding Values (EBVs). Active variants also revealed trait-related tissues, and single-cell RNA sequencing (scRNA-seq) identified the cell types most associated with production traits. To facilitate research, we developed the Integrated Functional Mutation (IFmut) platform, enabling users to explore variant functions easily. Our study provides a flexible platform and resource for studying genomic variation in farm animals, setting a new standard for research and breeding strategies.

**Conclusion:** The results indicated that evaluating functional potential by annotating and categorizing variants that interfere with transcription factor motifs can help elucidate changes in gene expression and phenotype. By focusing on high-confidence variants enriched in eQTL and trait-associated SNPs, it improves the accuracy of genomic predictions in research and breeding strategies.

## Introduction

Genomic variants in *cis*-regulatory elements are critical drivers of gene expression changes, influencing phenotypes [1-4]. While many variants are neutral, some can cause diseases or other harmful effects [5]. Identifying and characterizing variants is a critical focus in human genetics research, especially in understanding genetic disorders [6-8]. Large-scale initiatives like the 1000 Genomes Project have propelled human genome exploration [9, 10]. Similarly, the elucidation of genetic variants in farm animals, including pigs (*Sus scrofa*), cattle (*Bos taurus*), sheep (*Ovis aries*), chickens (*Gallus gallus*), mallards (*Anas platyrhynchos*), swan geese (*Anser cygnoides*), and goats (*Capra hircus*), as well as companion animals like rabbits (*Oryctolagus cuniculus*) and dogs (*Canis lupus familiaris*), and working animals like horses (*Equus asinus*), is vital for optimizing traits like production efficiency, health, behavior, and performance [11]. Genomic selection (GS) [12] has revolutionized animal breeding, enabling more precise selection based on traits like growth rates or disease resistance [13-17]. However, not all variants are functionally relevant, creating a need for systematic methods to identify and interpret functional variants.

We reviewed studies focused on identifying candidate functional variants. CADD, primarily designed for human variants, uses supervised machine learning to pinpoint deleterious mutations but may miss neutral or beneficial ones due to its bias [18]. Similarly, pCADD (for pigs) and chCADD (for chickens) follow similar algorithms but lack regulatory annotations, limiting their accuracy in non-coding regions [19, 20]. FitCons, which estimates selective pressure, may be less effective in detecting weakly selected mutations tied to complex traits [21]. Eigen offers robustness but relies on high-quality datasets, and the shortage of experimentally validated non-coding variant data hinders its reliability [22]. These methods struggle to predict non-coding variant effects in farm animals due to their reliance on conserved regions and limited data.

Functional genomic annotation is key to identifying variants associated with complex traits. Genome-wide association studies (GWAS) show that many genetic variants linked to diseases and traits lie in non-coding regions, as shown by the Encyclopedia of DNA Elements (ENCODE) [23]. In humans, the ENCODE project has driven considerable progress in functional variant screening, uncovering previously unrecognized functions of several regions in the human genome. In addition, RegulomeDB database and IGVF annotate and prioritize genetic variants in non-coding regions based on ENCODE project, enhancing the discovery of functional human variants and becoming a highly utilized resource for the research community [24, 25]. Despite these advances, regulatory variants in animal genomes remain underexplored.

To address the abovementioned concerns, we curated 877 datasets from 10 species, including pig, cattle, sheep, goat, horse, dog, rabbit, swan goose, mallard, and chicken. These include 357 ATAC-seq, 26 DNase-seq, 363 ChIP-seq, 56 Hi-C, 69 scRNA-seq, and quantitative trait loci (QTL) datasets related to trait of six species. Based on these, we annotated QTLs, open chromatin regions (OCRs), nucleosome-free regions (NFRs), motifs, footprints, TF-binding sites, and H3K27ac peaks, covering eight functional regulatory regions. We then identified candidate functional variants and developed a classification system to evaluate their impact on regulatory functions. We further assessed variant enrichment in expression QTLs (eQTLs) and trait-associated SNPs, the performance of high- and moderate-confidence variants in EBVs, variant activity across tissues, and integration of scRNA-seq data. Finally, we created an IFmut database and a Functional Confidence scoring system, providing a versatile public resource for researchers. This study delivers a comprehensive database and a powerful online toolkit, advancing the exploration of functional variants in farm animals for both research and breeding applications.

## Methods

### Data collection

Genome variants data in VCF format for pig (*Sus scrofa,* susScr11.1), cattle (*Bos taurus,* bosTau9), and chicken (*Gallus gallus,* galGal5) were downloaded from the Ensembl database (http://ftp.ensembl.org/pub/). Genome variants for mallard (*Anas platyrhynchos,* CAU_Wild1.0), goose (*Anser cygnoides*, GooseV1.0), goat (*Capra hircus*, ARS1.2), rabbit (*Oryctolagus cuniculus*, OryCun2.0), dog (*Canis lupus familiaris*, CanFam3.1), and horse (*Equus asinus*, EquCab3.0) were downloaded from Ianimal database (https://ianimal.pro/reception/download) [26], while sheep (*Ovis aries,* oviAri4) data were obtained from the NCBI Single Nucleotide Polymorphism Database (https://ftp.ncbi.nih.gov/snp/organisms/archive/sheep_9940/VCF/00-All.VCF.gz). QTL data for ten species were downloaded from the Animal QTL database (https://www.animalgenome.org/cgi-bin/QTLdb/). For pig, we also downloaded significant *cis*-eQTL and *trans*-eQTL data from the PigGTEx-Portal database (https://piggtex.ipiginc.com/#/downloads) [27].

TF ChIP-seq, H3K27ac ChIP-seq, ATAC-seq, and Hi-C data for pig, cattle, sheep, mallard, goat, goose, rabbit, dog, horse, and chicken, and scRNA-seq data in those eight species except goose and horse, as well as DNase-seq data for chicken, were downloaded from the NCBI Sequence Read Archive (http://www.ncbi.nlm.nih.gov/sra/). After quality control, a total of 802 epigenomic datasets and 69 scRNA-seq datasets were collected from multiple projects, including 75 datasets from pigs obtained from our previous study [28]. To investigate the evolutionary sequence conservation of functional variants, we downloaded GERP scores for pigs from Ensembl (https://ftp.ensembl.org/pub/current_compara/conservation_scores/).

### Sequencing data analysis

To adhere to ENCODE standards, we primarily followed the analysis methods outlined in our previous study for processing ChIP-seq and ATAC-seq data [28].

### ChIP-seq

#### Mapping and Quality control

The ENCODE ChIP-seq pipeline (https://github.com/kundajelab/chipseq_pipeline) was used to process ChIP-seq datasets. Raw reads from each dataset were aligned to the respective reference genomes using BWA v0.7.17 [29]. Low-quality reads (MAPQ < 25), unmapped reads, mate unmapped reads, non-primary alignment reads, and duplicate reads were removed using Picard v1.126 (https://broadinstitute.github.io/picard) and SAMTools v1.9 [30].

Read coverage of genomic regions between filtered replicate BAM files was computed using the multiBamSummary bins function of deepTools v2.0 [31], with a bin size of 2 kb to assess genome-wide similarities. Pearson correlation coefficients were calculated from the resulting read coverage matrix. Replicates with a Pearson correlation coefficient ≥0.8 were merged, while those with coefficients <0.8 were excluded from further analysis.

#### Identification of nucleosome-free regions

HOMER software was used to detect nucleosome-free regions (NFRs) [32]. Tag directories for H3K27ac IP and input data were generated using the makeTagDirectory command using the merged non-duplicated BAM file obtained from the “Mapping and Quality control” procedures. Subsequently, the findPeaks command with the -nfr option was applied to identify NFR peaks. Only datasets with at least 10,000 peaks were considered, excluding scaffold regions.

#### Identification of TF binding sites and H3K27ac narrow peaks

TF binding sites and H3K27ac narrow peaks were identified using MACS2 v2.1.0 [33] and deepTools v2.0 [31], as previously described in greater detail in our prior study [28].

### ATAC-seq

#### Mapping, quality control, and peak calling

The ENCODE ATAC-seq pipeline (https://github.com/kundajelab/atac_dnase_pipelines) was used to process ATAC-seq datasets. Adapter trimming was performed using Cutadapt v1.14 (https://cutadapt.readthedocs.io/en/stable/), followed by alignment to the reference genome assemblies with Bowtie2 v2.3.4.1. Low MAPQ reads (< 25), unmapped reads, mate unmapped reads, not primary alignments, reads failing platform, and duplicates were removed using SAMTools v1.9 [30] and Picard v1.126 (https://broadinstitute.github.io/picard). Mitochondrial reads were removed using BEDTools v2.26.0 [34] to generate effective reads for peak calling. Peaks were called using MACS2 v2.1.0 [33] using: genome size (-g), *p*-value threshold (0.01), peak model (--nomodel), shift size (--shift), extension size (--extsize), and other options (--B, --SPMR, --keep-dup all, --call-summits), to generate at least 10,000 peaks for further analysis.

### DNase-seq

#### Mapping, quality control, and peak calling

For the DNase-seq datasets for chicken, the ENCODE DNase-seq pipeline (https://github.com/kundajelab/atac_dnase_pipelines) was used, with ‘dnase_seq’ specified to indicate DNase-seq data. The remaining steps were consistent with the ATAC-seq analysis.

#### Identification of open chromatin regions (OCRs)

Peaks with *p* < 10^-5^ were considered significant for further analysis. Significant narrow peaks were filtered based on replicates with high Pearson correlation coefficients (*R* ≥ 0.8). Merged peaks, representing OCRs, were generated using BEDTools v2.26.0 [34], requiring at least 50% overlap between replicates. Signal tracks were generated from highly correlated replicates (*R* ≥ 0.8) using MACS2 v2.1.0 [33] to visualize chromatin accessibility across the genome.

#### Identification of footprints in ATAC-seq and DNase-seq

Footprint analysis followed these steps: (i) broad peaks were called from merged ATAC-seq or DNase-seq data using the MACS2 v2.1.0 broad module [33, 35]; (ii) significant broad peaks (*p* < 10^-10^ and 10^-10^ < *p* < 10^-5^ overlapping OCRs) were merged using BEDTools v2.26.0 [34]; (iii) Transcription factor footprinting was performed using the HMM-based Identification of Transcription factor footprints (HINT) framework from Regulatory Genomics Toolbox (RGT) v0.13.2 [36] with parameters adjusted for ATAC-seq or DNase-seq data (--atac-seq or --dnase-seq) and paired-end sequencing data (--paired-end). The organism information (--organism=) was also specified; and (iv) the cutoff for footprint score was set at the 20^th^ percentile of all footprints identified by the HINT framework of GRT v0.13.2 [36].

#### Transcription factor motif mapping in genome function region

The transcription factor motif mapping was primarily performed as follows: (i) OCR, NFR, TF binding sites, and footprint in OCR regions were merged into a BED file; (ii) The fasta-get-markov command from the MEME Suite (https://github.com/cinquin/MEME) was used to generate a .fa.bg file and the “bedtools getfasta” command generated the corresponding .fa file from the BED file of step (i); (iii) The fimo command in MEME Suite (--max-stored-scores 5,000,000) was used to map motifs in the genome; and (iv) the results of fimo mapping for pig, cattle, and sheep were filtered by *p* < 5×10^-6^, while for chicken, *p* < 5×10^-7^ was applied.

#### Prediction of transcription factor motif effects

Potential functional variants (Categories 1-5) located within footprint regions were analyzed using the motifbreakR [37] package in R v4.0. The motifDB database, specifically JASPAR 2018 [38], was selected as the data source for predicting the transcription factors to which the SNPs may bind.

#### Hi-C data processing

The Hi-C data of two-week-old LW pigs were taken from our previous study [28], and other Hi-C data were downloaded from the GEO at http://ncbi.nlm.nih.gov/geo. The downloaded data were processed using the HiC-Pro (v2.11.1) pipeline to produce the ICE normalization contact matrices [39]. The insulation score of the ICE matrix was calculated using the following options: -is 480000, -ids 320000, -im iqrMean, -ss 160000. The insulation method was utilized to define the topologically associating domain (TAD) structure (insulation/boundaries). Hi-C loops were analyzed using HiCCUPS v0.3.7 [40], with minor modifications, and the resolution parameter was set to “-r 25000, 40000”.

#### Analysis of scRNA-seq

A species-specific reference index was generated using “cellranger mkref” from 10x Genomics Cell Ranger v7.0.1. The following reference genomes and their corresponding filtered GTF files from Ensembl were used: pig (susScr11), cattle (bosTau9), chicken (galGal5), sheep (oviAri4), goat (ARS1.2), rabbit (OryCun2.0), and dog (CanFam3.1). The GTF files were filtered using “cellranger mkgtf” with default parameters. Expression data were processed using Seurat (v5.1.0) [41], with low-quality cells and empty droplets removed. Cells were initially filtered according to the following criteria: a minimum of 200 expressed genes per cell and at least three gene counts. Further quality control steps eliminated cells with fewer than 200 unique feature counts or mitochondrial content exceeding 5%. The remaining data were used for downstream analysis. Data normalization was performed using the NormalizeData function with a scaling factor of 10,000 to account for library sequencing depth. Principal component analysis (PCA) was conducted on the expression matrix of the top 2,000 most variable genes. Dimensionality reduction and clustering were performed using *t*-SNE. Cell types were identified using the CellMarker 2.0 database [42]. An R script was written to identify the intersection between the top 25 marker genes in each cluster and known marker genes from the database, enabling candidate cell type identification. The average log_2_FC of each marker gene was normalized, and the sum of all marker gene scores was utilized to assign a score to each cell type. Higher scores indicated a higher likelihood of the cluster representing a specific cell type. Final cell type identification was corroborated using prior research findings.

#### Variants distribution statistics

SNPs and small InDels were annotated using snpEff v4.5 software, with the reference genome file (FASTA) and annotation files (GTF) [43].

### Performance of genomic predictions

#### Dataset

The phenotypic dataset used for genomic prediction was obtained from a national pig nucleus herd in North China. The study included phenotypic recordings for two productive traits: average daily gain (ADG) from 30-100 kg and backfat thickness (BF) at 100 kg. All phenotypic records for the traits were obtained at the same time point, allowing a 10-kg deviation from the final body weight (100 ± 10 kg). All phenotypes were recorded between early 2018 and October 2022. Based on the traced pedigree, 11 pig lines were identified within the population. DNA samples were collected from approximately 80 distantly related pigs per line, and sequencing was performed using the DNBSEQ-T7 platform, with an average coverage of 5×. In total, 874 pigs were sequenced. After quality controls–such as a genotype missing rate below 10%, a call rate of SNPs above 90%, and a minimum allele frequency (MAF) above 1%–18,460,807 (18000K) SNPs were retained for analysis. Missing genotypes were imputed using Beagle software (v5.3). Of the 874 sequenced pigs, 872 had ADG phenotypic data, while 867 had BF recordings. Environmental factors, including gender, herds, and physical units, were fully recorded.

#### Genomic Best Linear Unbiased Prediction (GLUP) models

Breeding values (EBV) for different traits were estimated using GBLUP models:

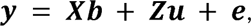

where ***y*** indicates a column vector of phenotypic values for each trait; ***b*** represents a vector of fixed effects, including sex effect, herd effect, and physical unit effects; ***u*** denotes a vector of random additive genetic effects; ***e*** is a vector of residual effects. Matrices ***X*** and ***Z*** are design matrices associated with these effects. The GBLUP model assumes a normal distribution for the random additive and residual effects, as 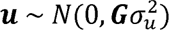, where G denotes a genomic relationship matrix constructed as Vanraden method 1; 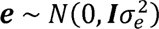, where I is an identity matrix. The additive genetic variance and residual variance are indicated by 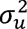 and 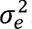, respectively.

#### Scenarios of constructing genomic relationship matrices

GBLUP models with four different genomic relationship matrices (***G***) were employed to estimate GEBVs for ADG and BF traits. Four different sets of SNP markers were utilized to develop corresponding ***G*** matrix. In scenario 1, sequenced SNP markers with the top 1 and top 2 muscle scores (1+2 muscle, 10,544 SNPs) were calculated the ***G*** matrix. Similarly, sequenced SNP markers with the top 1 and top 2 liver scores (1+2 liver, 6,049 SNPs) and top 1 and top 2 adipose scores (1+2 adipose, 3,801 SNPs) were employed to construct ***G*** matrices in scenarios 2 and 3, respectively. In scenario 4,11,000 randomly selected SNP markers were utilized to develop a G matrix. Scenario 4 was repeated three times in the study.

#### Predictive Reliabilities

The mean predictive reliabilities of GEBVs were computed using the subsequent formula [44]:

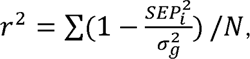

where *r*^2^ is the reliability of GEBVs and *i* indicates an individual animal i; SEP denotes the standard error associated with the predicted GEBVs; 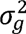 is the additive genetic variance and *N* is the number of animals used.

## Results

### Large-scale epigenomic screening of functional variants across farm animals

To annotate genomic variation, we gathered SNPs and small InDels (2-50 bp) from public databases for ten farm animals: pig, cattle, sheep, goat, chicken, horse, mallard, swan goose, rabbit, and dog. This dataset, comprising nearly 570 million variants, was classified using various sequencing methods (Fig. 1A). SnpEff software analysis showed that over 95% of the variants were located in non-coding regions across all species (Fig. 1B), consistent with previous findings in human studies where variants are commonly found in intergenic and intronic regions [7, 45, 46]. Thus, identifying functional variants from this vast pool remains a significant challenge. Given that regulatory regions, particularly transcription factor (TF) binding sites identified by epigenomic analyses, can inform the potential functions of variants, we aimed to screen for potentially functional variants using epigenomic data [25, 47]. To do this, we collected datasets from previous studies, including the FANNG project [48, 49] and our own research [28], encompassing ATAC-seq, DNase-seq, H3K27ac ChIP-seq, TF ChIP-seq, Hi-C, and scRNA-seq data from 10 farm animals. After rigorous mapping and quality control, 871 datasets from 32 tissues and eight cell lines passed quality standards, including 357 ATAC-seq, 26 DNase-seq, 363 ChIP-seq, 56 Hi-C, and 69 scRNA-seq datasets (Fig. 1C). Specifically, we compiled 160, 199, 40, 22, 106, 213, 11, 8, 31, and 81 datasets for pig, cattle, sheep, goat, horse, chicken, mallard, swan goose, rabbit, and dog, respectively.

**Figure 1.**
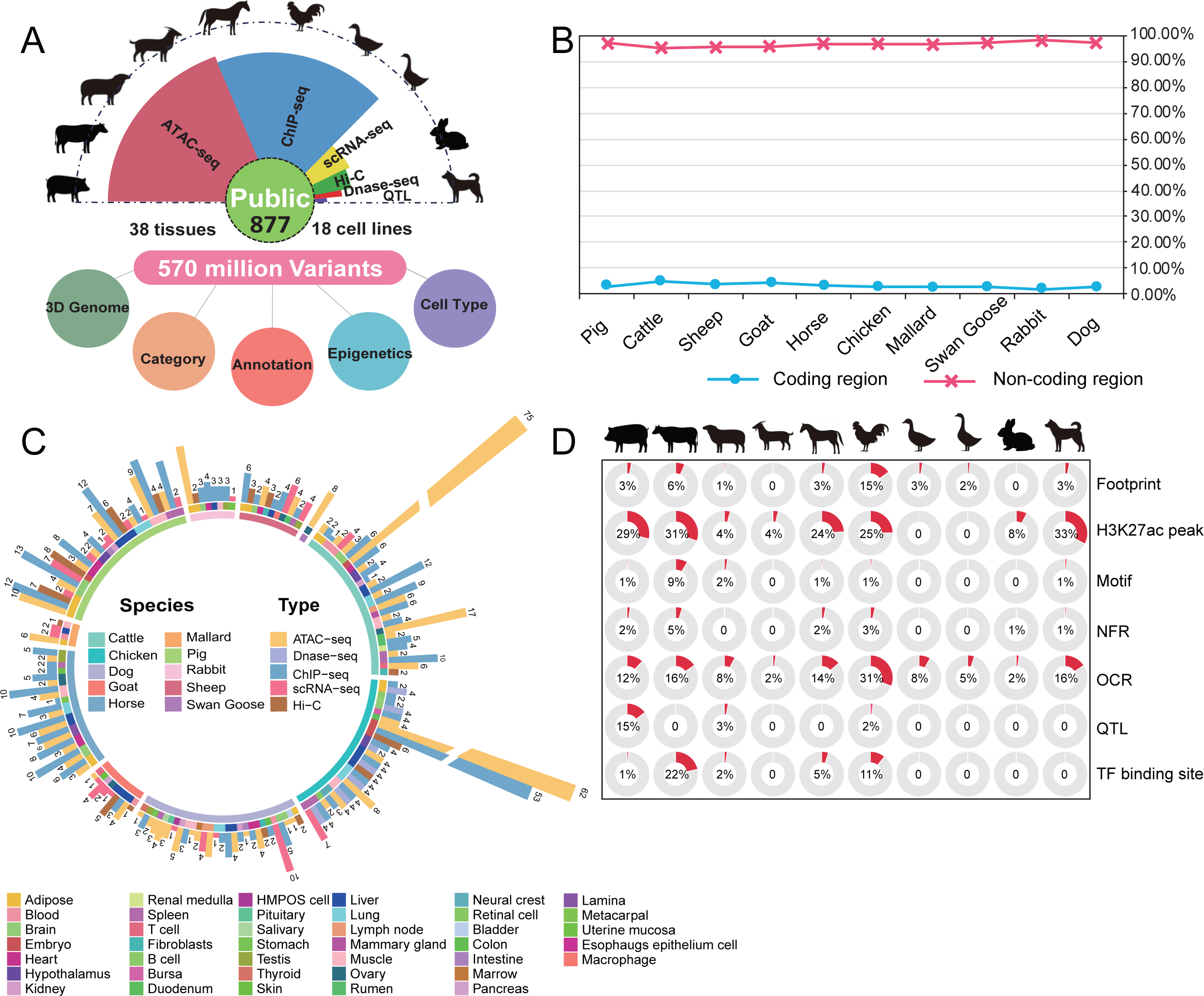
Workflow and genomic distribution of variants and epigenomic datasets collection across ten farm animals. (A) Overview of the workflow. (B) Proportion of variants distributed across coding and non-coding regions in the ten species. (C) Statistical summary of datasets categorized by tissue types across the ten species. The color of the inner circle represents the species, the middle circle represents tissues and cell lines, and the outer circle indicates the number of datasets corresponding to different data types. (D) The proportion of genomic variants within different functional regions of the ten species.

Using ENCODE guidelines (https://www.encodeproject.org/), we identified regulatory regions, resulting in over 4.6 million non-redundant genomic regions across all species. These included open chromatin regions (OCR), H3K27ac peaks, footprints, and nucleosome-free regions (NFR) (Supplementary Table S1). Additionally, TF binding peak calling combined with positional weight matrices (PWMs) identified 8.8 million non-redundant regions associated with TF binding sites across the ten species (Supplementary Table S1). The total length of these regions accounted for 33.55% of the pig genome (Sscrofa11.1), 43.67% for cattle (ARS-UCD1.2), 15.00% for sheep (Oar_v4.0), 5.12% for goat (ARS1.2), 34.76% for horse (EquCab3.0), 44.65% for chicken (Gallus_gallus-5.0), 7.94% for mallard (CAU_Wild1.0), 4.37% for swan goose (Goose1.0), 8.93% for rabbit (OryCun2.0), and 38.97% for dog (CanFam3.1) (Supplementary Table S2).

Genomic variants in TF binding sites, such as those cataloged in RegulomeDB, often result in functional consequences [25]. Since the above genomic regions containing basic regulatory and TF binding-related regions were identified through DNA-TF interaction data, variants detected in these regions were likely to have transcription regulation function in the farm animals. Moreover, we further identified 1,128,759∼39,157,953 (4.98∼47.05%) variants located in these regulatory regions across the ten species (Supplementary Fig. S1), which we designated as potential functional variants.

### Classification approach for functional impact of variants

Gene expression is largely regulated by TFs, proteins that bind to specific [50] DNA sequences to control gene transcription [50, 51]. Variants located within TF binding motifs can disrupt this binding, potentially altering gene expression and impacting phenotype (Fig. 2A). To better evaluate the likelihood that a regulatory variant impacts transcription, we developed a functional confidence classification system similar to that of RegulomeDB [25] (Fig. 2B; Supplementary Table S3).

**Figure 2.**
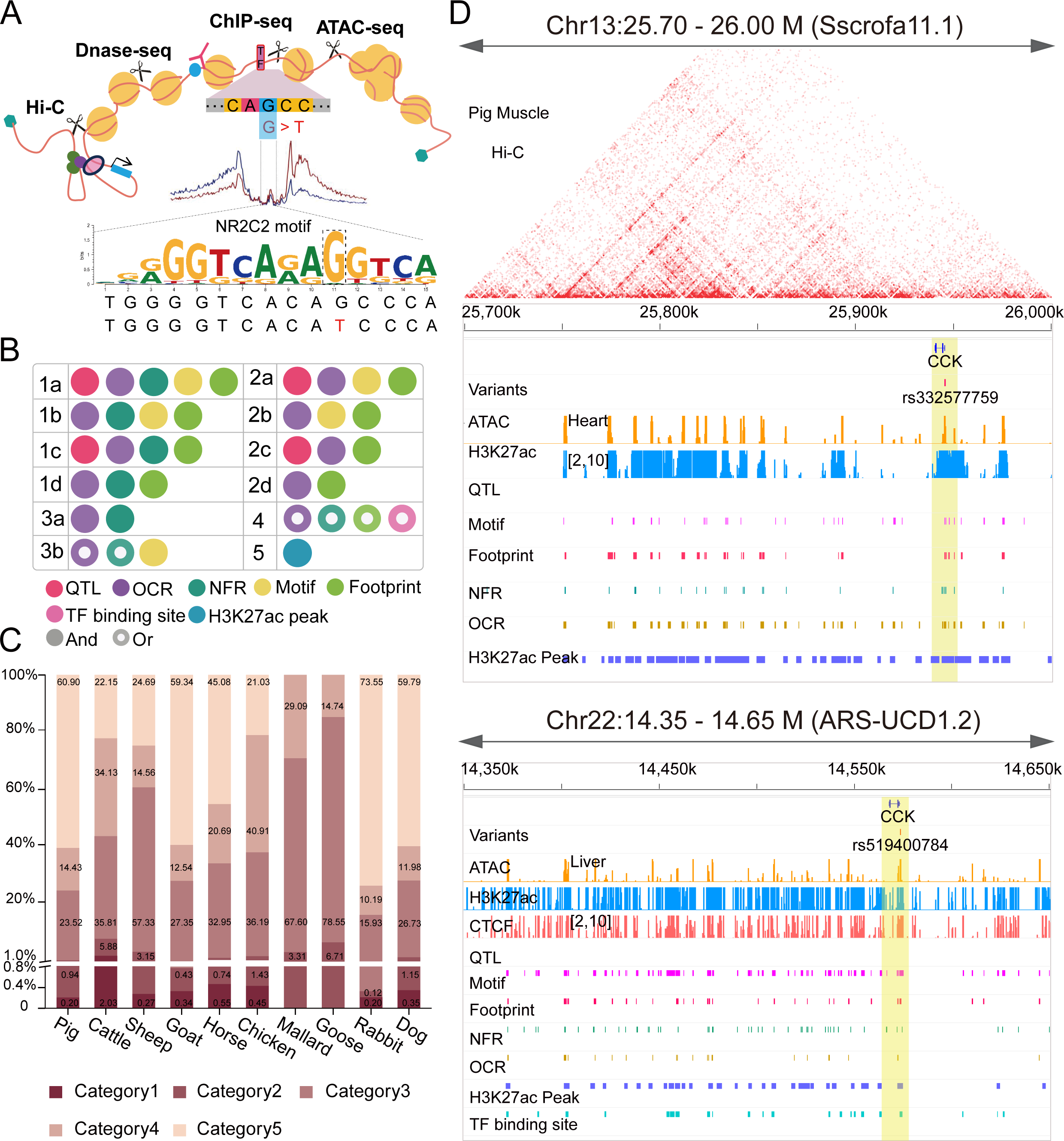
Functional classification of genomic variants: approach and findings. (A) Schematic diagram illustrating how genomic variants influence transcription factor binding. (B) Design principles underlying the system for classifying variant functional confidence. (C) Proportions of variations from categories 1-5 across the ten species. (D) Signal distributions surrounding variants within the *CCK* gene in pigs and cattle. The top panel shows ATAC-seq and H3K27ac ChIP-seq signal peaks around the SNP rs332577759 (Category 2d) in heart tissue, alongside functional regions and interaction heatmaps in muscle. The lower panel displays rs519400784 (Category 1d) signal intensity in cattle liver, with key functional regions highlighted in yellow.

Currently, only 103 TF ChIP-seq datasets exist for farm animals, most of which target CTCF (86 datasets). Chickens have the largest number of ChIP-seq datasets, with 17 datasets for 9 TFs. Thus, due to the lack of TF ChIP-seq data in farm animals, the identification of potential TF motif regions relied on a combination of ATAC-seq/Dnase-seq (i.e., OCR and footprints) and H3K27ac ChIP-seq (i.e., NFR and H3K27ac peaks). We also incorporated QTL data, as variants in these regions may affect agronomic traits.

Our functional confidence classification system evaluates variants based on the number of regulatory regions they overlap with. Given the prominent association of NFRs, OCRs and TF footprints (especially those containing fully or partially matching recognition motifs) with transcriptional activation, variants in these regions had the highest likelihood of affecting TF binding and gene expression, and were therefore scored as high functional confidence variants (Category 1). Variants meeting these criteria but never found in NFRs were subsequently scored as moderate functional confidence (Categories 2a-2d), suggesting a moderate likelihood of affecting TF activity. Additionally, variants located in QTL regions were assigned higher scores (e.g., Categories 1a, 1c, 2a, and 2c), while those outside QTLs received slightly lower scores (Categories 1b, 1d, 2b, and 2d). By contrast, variants detected in DNase/ATAC-seq or H3K27ac ChIP-seq data but not in TF footprints or recognition motifs were classified as low (Category 3) or minimal (Category 4) functional confidence. Category 5 included variants detected only by H3K27ac ChIP-seq, which may still influence transcription regulation (Fig. 2B; Supplementary Table S3).

Based on our classification approach, we identified 126,937,925 candidate functional variants. Among these, 26,341 to 3,096,314 SNPs/small InDels were classified as high or moderate confidence (Categories 1 and 2), representing 1.15% to 7.91% of all functional variants across the ten animals (Fig. 3C; Supplementary Fig. S2). As expected, we observed some variants with functional potential were related to genes known to significantly influence production traits. For example, the CCK gene encodes cholecystokinin, which regulates digestive processes, appetite, and fat metabolism, playing a crucial physiological role in animals [52-55]. In pig genome, there is a SNP (rs332577759, chr13:25945714:C:T), which located in the exon of CCK, showed significant signal in the ATAC-seq and H3K27ac ChIP-seq peaks, leading to its annotation and classification as Category 2d (Fig. 2D). Similarly, the SNP rs519400784 (chr22:14573586:C:CCCGAGAG) from cattle genome, located in the homologous region of rs332577759, is also surrounded by strong signals of ATAC-seqH3K27ac ChIP-seq, and CTCF ChIP-seq, resulting to its classification as Category 1d (Fig. 2D).

**Figure 3.**
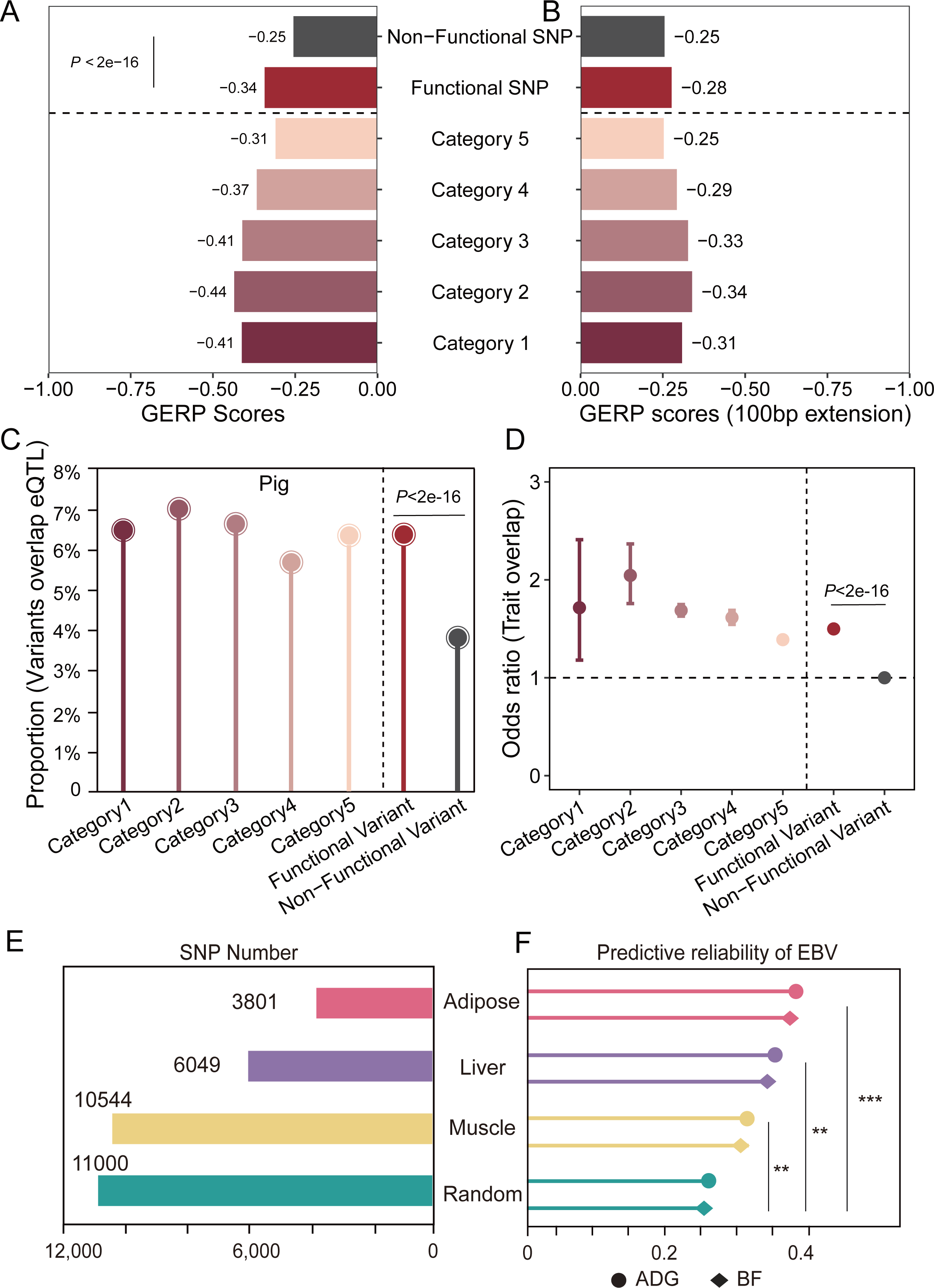
Conservation and impact of functionally significant variants. (A) GERP scores (from Ensembl) for variants across different functional categories in the pig genome. (B) GERP scores for the 100 bp regions flanking the variants (bedtools slop -b 100). (C) Proportion of variants overlapping with eQTLs. (D) Fisher’s exact test odds ratios for variants linked to production traits (data from Animal QTL), with non-functional variants serving as the control. (E) Number of SNP markers used to evaluate EBV. (F) Reliability of EBV predictions for traits like ADG and BF.

### Variants in higher functional categories are more likely associated with traits

To further explore the characteristics of variants across different functional categories, we utilized the Ensembl Genomic Evolutionary Rate Profiling (GERP) score to assess the evolutionary conservation of all variants [56]. Surprisingly, variants with higher functional confidence showed significantly lower GERP scores compared to non-functional ones, especially in Category 1 (Fig. 3A). Negative GERP scores indicate reduced sequence conservation at these loci, suggesting higher mutation rates, potentially due to neutral or positive selection. To ensure that these conservation differences weren’t influenced by the surrounding regulatory elements, we extended the analysis to include 100 bp flanking regions around the variants (Fig. 3B). The results showed that GERP scores in non-functional variants remained stable, while the conservation scores of functional confidence variants increased significantly, approaching the levels observed in non-functional regions. This finding supports the notion that the lack of conservation is an inherent feature of these variants.

Previous studies suggest that artificial selection has shaped farm animal genomes, particularly in loci linked to production traits, which tend to undergo strong positive selection [57]. This evolutionary pattern mirrors what we observed in variants with higher functional categories, leading us to hypothesize that artificial selection may drive the lower conservation of these variants. To test this, we analyzed eQTL data from over 30 pig tissues using the PigGTEx database and observed that eQTLs were significantly more likely to be functional variants than non-functional ones (Fig. 3C) [27]. This indicates that functional variants are more likely to influence gene expression, a key determinant of phenotypic traits. Additionally, intersecting functional confidence variants with known trait-associated loci revealed significant enrichment in these regions (Fig. 3D). These findings demonstrate that our Functional Confidence Classification system effectively identifies variants with a high likelihood of impacting gene expression and driving phenotypic variation.

To further validate our classification system, we screened high and moderate confidence variants (Category 1 and Category 2) in pigs and assessed their predictive reliability for estimated breeding values (EBVs) in two traits—average daily gain (ADG) and backfat thickness (BF)—using a large white population (*n* = 874). Genomic BLUP modeling was performed using DMU software [58] with four scenarios: three using functional variants from muscle, liver, or adipose tissue, and one using 11K random variants from whole-genome pig sequencing (Fig. 3E; Supplementary Table S4). The predictive reliability of EBVs for ADG and BF was similar across the three tissue-based scenarios (∼0.31–0.38), with the adipose tissue scenario showing the highest reliability (∼0.38), despite using the fewest markers (3,861 SNPs). In comparison, the random SNP scenario, despite containing the highest number of markers (11,000 SNPs), yielded the lowest reliability (∼0.27). The predictive reliability of EBVs improved by 23–46% for all three tissue types when high and moderate functional variants were used, compared to the random SNP scenario (Fig. 3F). This further demonstrates the utility of our Functional Confidence Classification system in screening functional variants for genomic data in farm animals.

### Functional variants define the principal tissues of *cis*-regulatory elements and traits

Besides evaluating the functional potential of variants, considering that the function of variants is heavily influenced by their regulatory elements, our data also enables the identification of the specific tissues in which these variants are active. By merging regulatory elements, we generated a comprehensive *cis*-regulatory element (CRE) map for seven species (the tissue numbers in other species were insufficient). The proportion of active variants was used to estimate the tissue-specific activity of CREs, and as expected, most CREs exhibited strong tissue specificity, being active in only one or two tissues (Fig. 4A; Supplementary Fig. S4). This observation aligns with findings from the human EpiMap project [59], which reported that 88% of enhancer elements are tissue-specific. Our results further extend this conclusion, showing that such tissue specificity is prevalent across all types of CREs.

**Figure 4.**
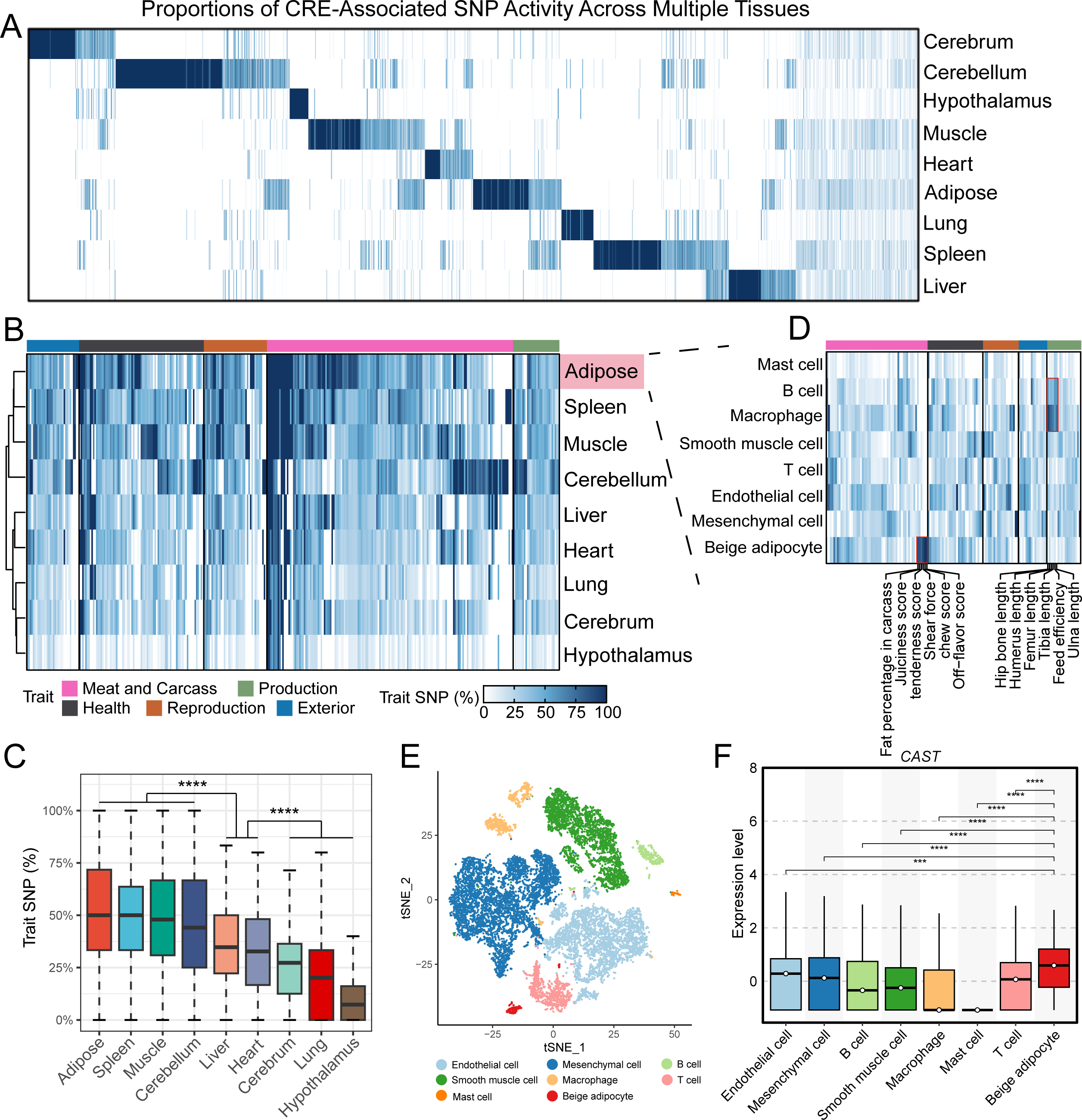
Variant activity in CRE and traits. (A) Proportions of CRE-linked SNP activity across various pig tissues. (B) Heatmap showing the enrichment of production traits based on the activity of trait-associated variants across pig tissues. (C) Proportion of active variants linked to specific traits. Statistical significance was determined using an unpaired Student’s *t*-test. * *p* < 0.05, ** *p* < 0.01, *** *p* < 0.001, **** *p* < 0.0001. (D) Enrichment of production traits across all cell types within adipose. (E) T-distributed stochastic neighbor embedding (t-SNE) plot showing scRNA-seq data from pig adipose tissue. (F) Expression levels of the *CAST* gene across various cell types.

Next, we analyzed the activity of trait-associated variants to identify the tissues linked to production traits. In animal breeding, accurate identification of tissues related to production traits is critical for understanding the underlying regulatory mechanisms and improving breeding strategies [60, 61]. To this end, we quantified trait-associated variants in each tissue and calculated their proportion relative to all variants related to the trait. Hierarchical clustering grouped tissues into three distinct clusters (Fig. 4B). Tissues such as adipose, spleen, muscle, and cerebellum showed significant enrichment of trait-associated variants, indicating a strong association with production traits (Fig. 4C). In contrast, tissues like the lung, cerebrum, and hypothalamus showed considerably lower proportions of trait-associated variants (Fig. 4C).

To further delineate the cellular origins of these production traits, we employed single-cell RNA-Seq (scRNA-seq) data to identify the cell types where variant-related genes were classified as marker genes (Fig. 4D and 4E). Variants were considered active in a cell type when their associated gene was a marker gene, reflecting the functional relevance of the variant in that cellular context. Through calculating the proportion of active variants across all traits, we observed some traits related to skeletal growth, such as feed efficiency and bone lengths (hip, humerus, femur, tibia, and ulna), were enriched in B cells and macrophages, highlighting the importance of immune cells in skeletal development (Fig. 4D). This finding aligns with recent research highlighting the interaction between the immune and skeletal systems, where immune cells regulate bone remodeling via cytokine, chemokine, and growth factor secretion [62, 63].

Moreover, several traits related to flavor and texture, such as fat percentage in the carcass, juiciness, tenderness, shear force, chew score, and off-flavor score, were significantly enriched in beige adipocytes. These traits, involving 33 SNPs, were nearly all associated with the *CAST* gene (Fig. 4F). We found that *CAST* expression was significantly higher in beige adipocytes than in other cell types. Previous studies have shown that the *CAST* gene is linked to meat tenderness, muscle growth rate, and post-mortem proteolysis [64], which are key factors influencing pork quality. Our findings align with these studies and further demonstrate that the *CAST* gene’s influence on flavor-related traits is specifically localized to beige adipocytes.

### Development of the Integrated Functional Mutation (IFmut) database for screening candidate functional variants in farm animals

In order to facilitate screening for candidate functional variants in farm animals, we integrated genomic variants with epigenomic datasets in a database, the Integrated Functional Mutation (IFmut) database. IFmut (http://www.ifmutants.com:8212) is a comprehensive farm animal variants database which provides functional context to variants or regions of interest and serves as a tool to prioritize functionally important variants (Fig. 5A). There are six modules (Category, Epigenetics, 3D Genome, Gene, Trait, and Single cell) within IFmut that enable users to query variants of interest (Fig. 5B).

**Figure 5.**
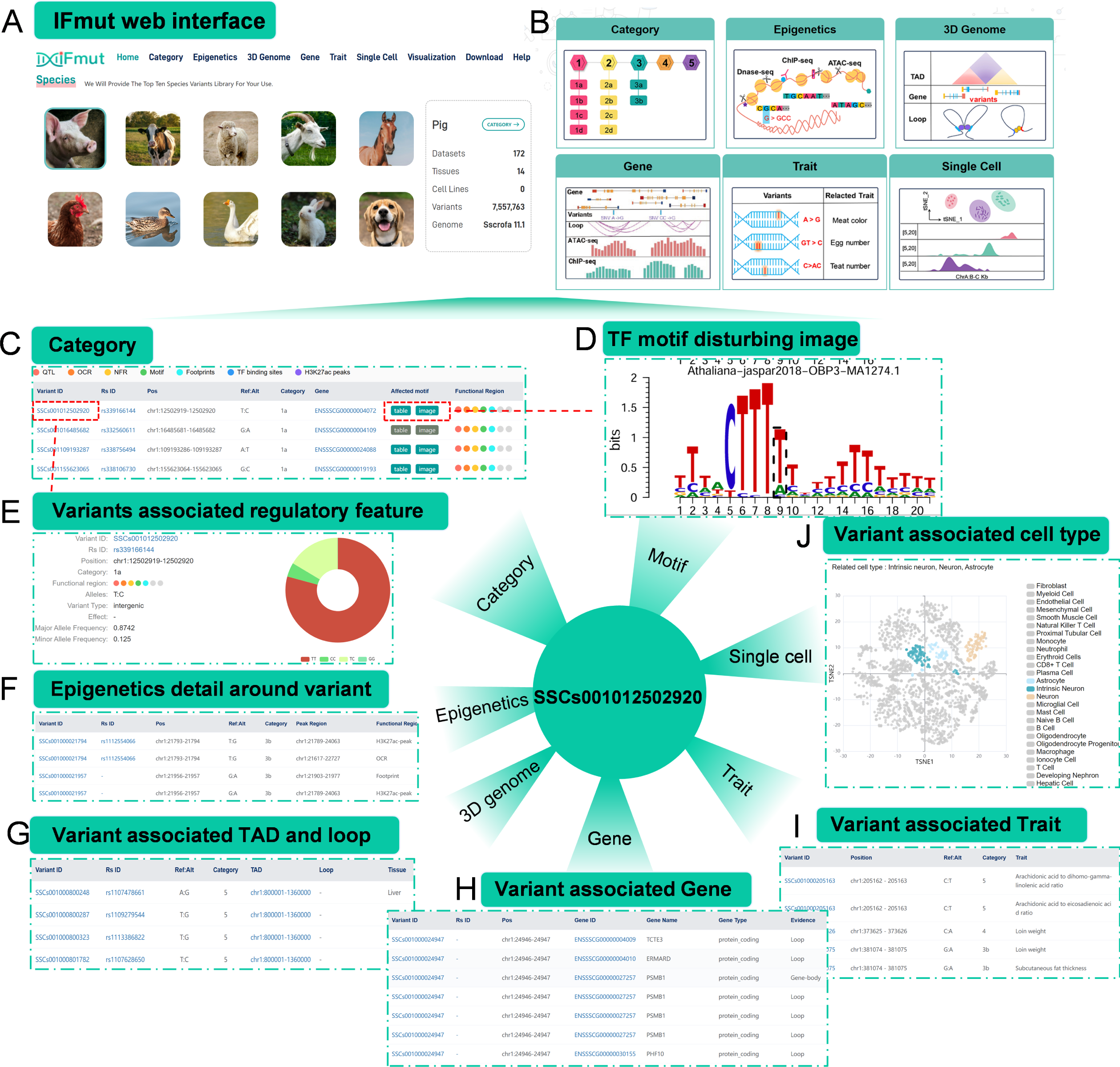
Overview of the IFmut database. (A) The IFmut homepage, displaying information about the ten farm animals featured. (B) Graphic overview of the modules available within IFmut. (C) Example of the variant category browser on the “Category” page. (D) Clicking the “image” button in the “Affected motif” column displays nucleotide conservation logos for TF recognition motifs potentially disrupted by a variant. (E) The variant ID link redirects users to detailed variant information. (F) Example of the epigenetics data browser for variants, available on the “Epigenetics” page. (G) Browser showing TADs or loops encompassing the variant on the “3D genome” page. (H) Browser the variant-related genes on the “Gene” page. (I) Trait browser displaying traits associated with the variant on the “Trait” page. (J) Cell type browser showing cell types linked to the variant on the “Single cell” page.

The ‘Category’ module integrates functional confidence categories for variants from ten species, facilitating searches by variant ID or position (Fig. 5C). Additionally, we also provide a tool (IfmutAnnotator) that allows users to upload a BED file containing variants not included in our database to classify novel variants (Supplementary Fig. S5). The “Motif” column aids in hypothesizing the variant’s potential impact on TF binding (Fig. 5D). Users can access detailed annotations by clicking on the variant ID (Fig. 5E). The ‘Epigenetics’ module offers six types of epigenetic regulatory regions (OCR, NFR, Footprint, Motif, TF-binding site, and H3K27ac peak) across tissues or cells (Fig. 5F), and the ‘3D Genome’ module contains interaction data such as loops and TADs (Fig. 5G). In the ‘Gene’ module, variants are linked to genes within the same TAD or chromatin loop, or to the nearest functional gene when 3D genome data is unavailable (Fig. 5H). The ‘Trait’ module enables users to search for trait-associated variants, providing insights into the genetic mechanisms of complex traits (Fig. 5I), while the ‘Single Cell’ module uses curated scRNA-seq data to analyze variant-associated gene expression across cell types (Fig. 5J). Visualization tools allow users to view epigenetic and Hi-C interaction signals around variants and nearby genes.

In summary, IFmut offers seven key types of functional information and practical tools, supporting the exploration of variant functionality.

## Discussion

Previous research has revealed that most SNPs and small InDels are located in non-protein-coding regions of the genome [65-68], making it challenging to interpret how these variants affect function [69, 70]. Evaluating the impact of variants on TF binding sites is an effective approach for identifying functional variants in the human genome [25], but the lack of TF ChIP-seq data for farm animals has hindered this method’s application in livestock research. To overcome this limitation, we utilized alternative epigenomic techniques like ATAC-seq and DNase-seq, which capture the accessible chromatin landscape and provide insights into TF binding sites. These techniques have been widely used in human and livestock research [71-73].

By leveraging epigenomic datasets, we annotated regulatory regions and developed a confidence classification system for variants based on their functional relevance. Overall, our approach was more suitable the current study of functional variants in livestock for abundant epigenetic datasets in these species, as well as the design idea of using the epigenetic data to identify TF binding sites to rank functional variants can also be transplanted to related research works on other species.

Moreover, by integrating eQTL data and trait-associated variants with our classification system, we demonstrated that higher functional confidence variants are more likely to influence gene regulation and trait expression. This enhances the practical utility of our system. For example, genomic predictions based on high- and moderate-confidence variants improved the reliability of Estimated Breeding Values (EBVs) in pigs compared to predictions using randomly selected SNPs. Furthermore, our analysis of active variants allowed us to identify the tissues associated with key production traits, such as adipose, spleen, and muscle tissues, aligning with breeding insights. Additionally, by incorporating scRNA-seq data, we pinpointed specific cell types contributing to these traits. These findings suggest that our functional classification system can effectively guide the identification of critical variants and thus contribute to advancing genetic improvement in farm animals.

The lack of research into regulatory variants in farm animal genomes has constrained progress in animal genomics, genetics, and breeding research. To address this gap, we designed the IFmut webserver, which enables users to explore genomic and epigenetic data related to variant functionality. Unlike other resources such as PigBiobank, AGIDB, and IAnimal [26, 74, 75], IFmut considered the regulation of genomic interactions that may result from SNP interference with TF. We also incorporated Hi-C data to characterize 3D genomic interactions and utilized single-cell data to assess the specificity of variants across cell types. Additionally, IFmut includes a Functional Confidence classification tool for evaluating novel variants, which is essential for advancing our understanding of how genetic variants influence traits, ultimately supporting more informed decision-making in livestock breeding and genetic improvement.

## Conclusion

Assessing functional potentiality by annotating and classifying variants that disrupt transcription factor motifs can explain alterations in gene expression and phenotype. Variants with high confidence were enriched in eQTL and trait-associated SNPs, show greater potential to influence gene expression and phenotype and help improve the accuracy of EBVs. The establishment of the IFmut platform provides a flexible platform and resource that sets a new standard for research and breeding strategies.

## Abbreviations

TF: Transcription factor
EBVs: Estimated Breeding Values
scRNA-seq: Single-cell RNA sequencing
IFmut: Integrated Functional Mutation
GS: Genomic selection
GWAS: Genome-wide association studies
ENCODE: Encyclopedia of DNA Elements
QTL: Quantitative trait loci
OCRs: Open chromatin regions
NFRs: Nucleosome-free regions
PWMs: Positional weight matrices
GERP: Ensembl Genomic Evolutionary Rate Profiling
ADG: Average daily gain
BF: Backfat thickness
CRE: *Cis*-regulatory element
PCA: Principal component analysis
GLUP: Genomic Best Linear Unbiased Prediction

## Acknowledgement

We thank Dr. Christopher K Tuggle at Department of Animal Science, Iowa State University for valuable suggestions to our work.

## Author contributions

Y.Z, T.X, and M.S conceived and designed the project. R.M, R.K, J.Z, J.S, Y.X, X.Z, Z.H, M.H, D.W, Y.F, Y.Z, X.L, M.Z and S.Z performed the analysis. M.S and Y.F constructed the web-based resource with input from R.M and M.H. R.M, R.K, J.Z, and M.S wrote the manuscript and prepared the figures. All authors read and approved the final manuscript.

## Funding

This work was supported by the National Natural Science Foundation of China (32341051), the grant from Department of Agriculture and Rural Affairs of Hubei Province (HBZY2023B006-02), National Key R&D Young Scientists Project (2022YFD1302000), Fund of Modern Industrial Technology System of Pig (CARS-35).

## Availability of data and materials

All ATAC-seq, ChIP-seq (H3K27ac and TFs), and Hi-C, as well as the candidate functional variants and their Functional Confidence scores are available at http://www.ifmutants.com:8212. The source code for calculating the Functional Confidence score for candidate functional variants is publicly available on GitHub at https://github.com/RXMa-Bio-software/the-Functional-Confidence-scoring-system.

## Declarations

### Ethics approval and consent to participate

Not applicable.

### Consent for publication

Not applicable.

### Competing interests

The authors have no competing interests to declare.

## Supporting information

**Supplementary Table S1. Number of regulatory regions detected in farm animals.**

**Supplementary Table S2. Genomic coverage of regulatory regions in farm animals.**

**Supplementary Table S3. Variant classification approach.**

**Supplementary Table S4. Predictive reliabilities of estimated breeding values for traits ADG and BF.**

**Supplementary Fig. S1. Number of variants in different functional regions of 10 species.** Including variants in the functional regions of motif, TF binding site, NFR, footprint, OCR, H3K27ac peaks and the number of variants not in any of the functional regions.

**Supplementary Fig. S2. Statistics on the number of variants in categories 1-5 (12 subclasses) for 10 species.**

**Supplementary Fig. S3. Conservation calculations of functional variants and non-functional variants for 6 species.** There were significant differences in the conservatism of functional variants and non-functional variants.

**Supplementary Fig. S4. Conservation calculations of functional variants and non-functional variants for 6 species.** There were significant differences in the conservatism of functional variants and non-functional variants.

**Supplementary Fig. S5. Instructions for using the ifmutAnnotator tools in IFmut.** On the Category page of the IFmut database, we provide the ifmutAnnotator tools annotation noval variant. The use of the ifmutAnnotator tool is divided into 3 steps, step1, click on the browse button, step2, upload the bed file to be annotated, and step3, download the annotation results.

## Notes

http://www.ifmutants.com:8212

### Competing Interest Statement

The authors have declared no competing interest.

### Summary of Updates

We have updated all the Supplemental files,supplementary explanation to the previous content.

