## Supplementary figures and images for "Annotation and assessment of functional variants in regulatory regions using epigenomic data in farm animals"

### Supplementary Figure 1

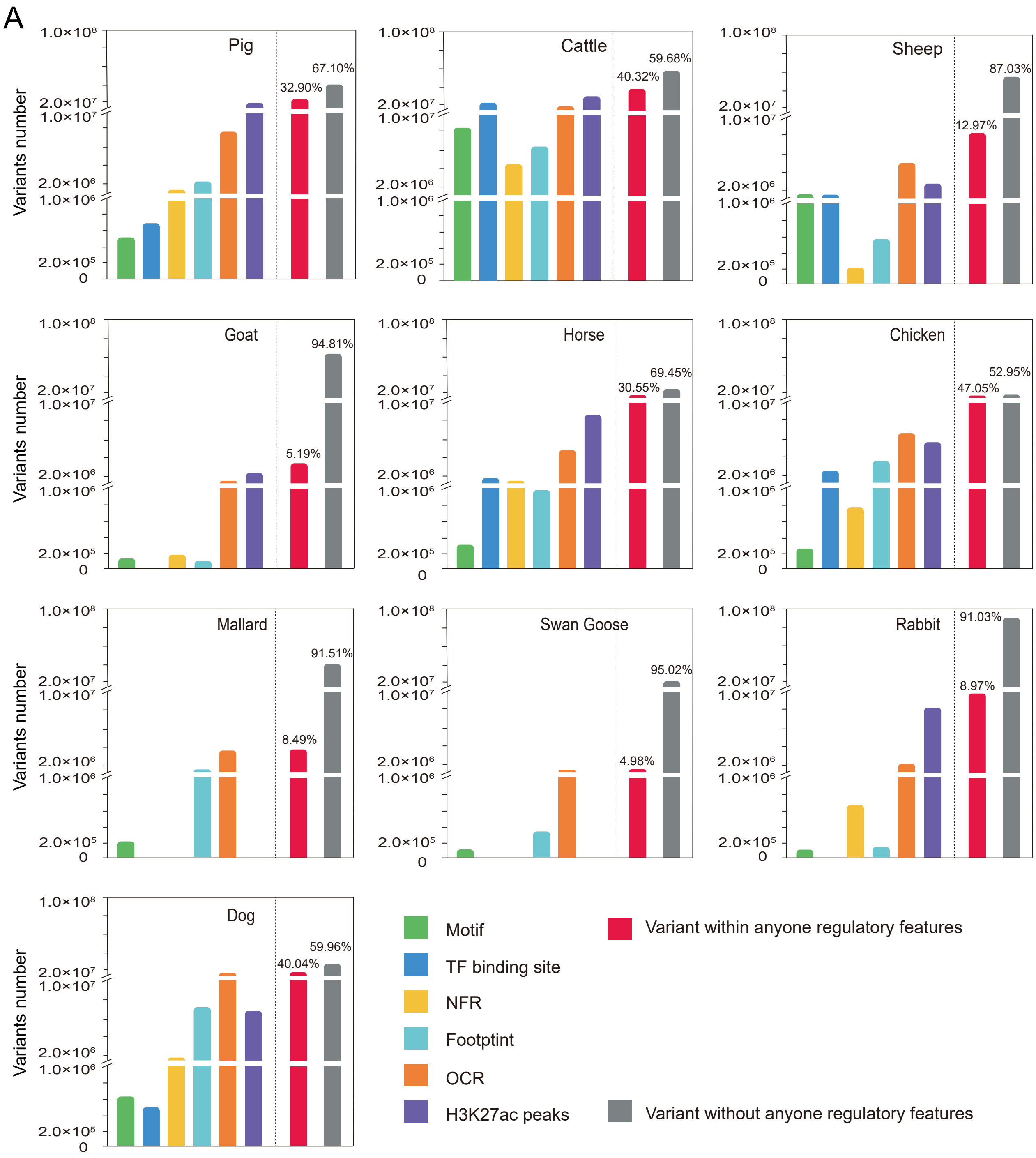

### Supplementary Figure 2

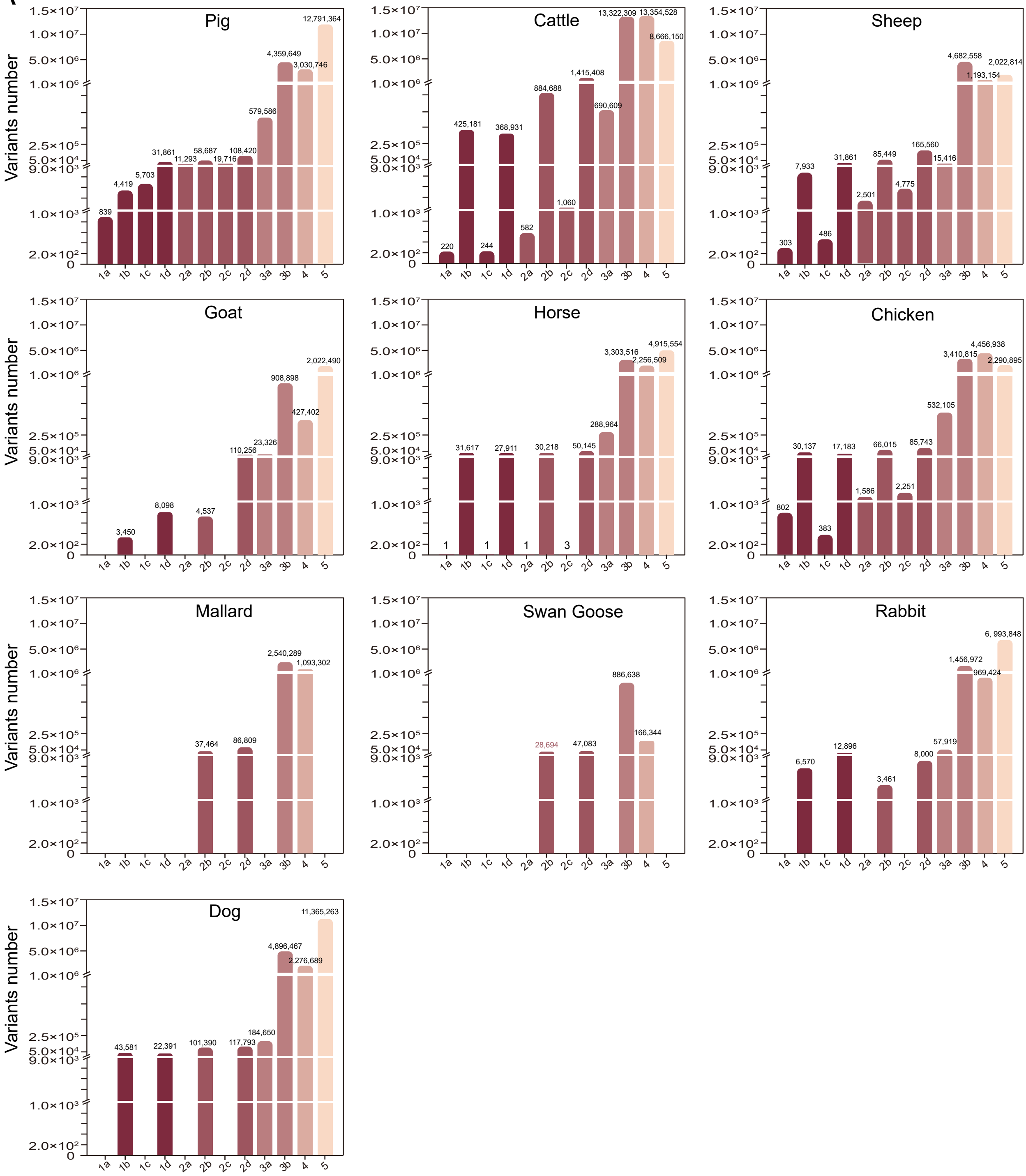

### Supplementary Figure 3

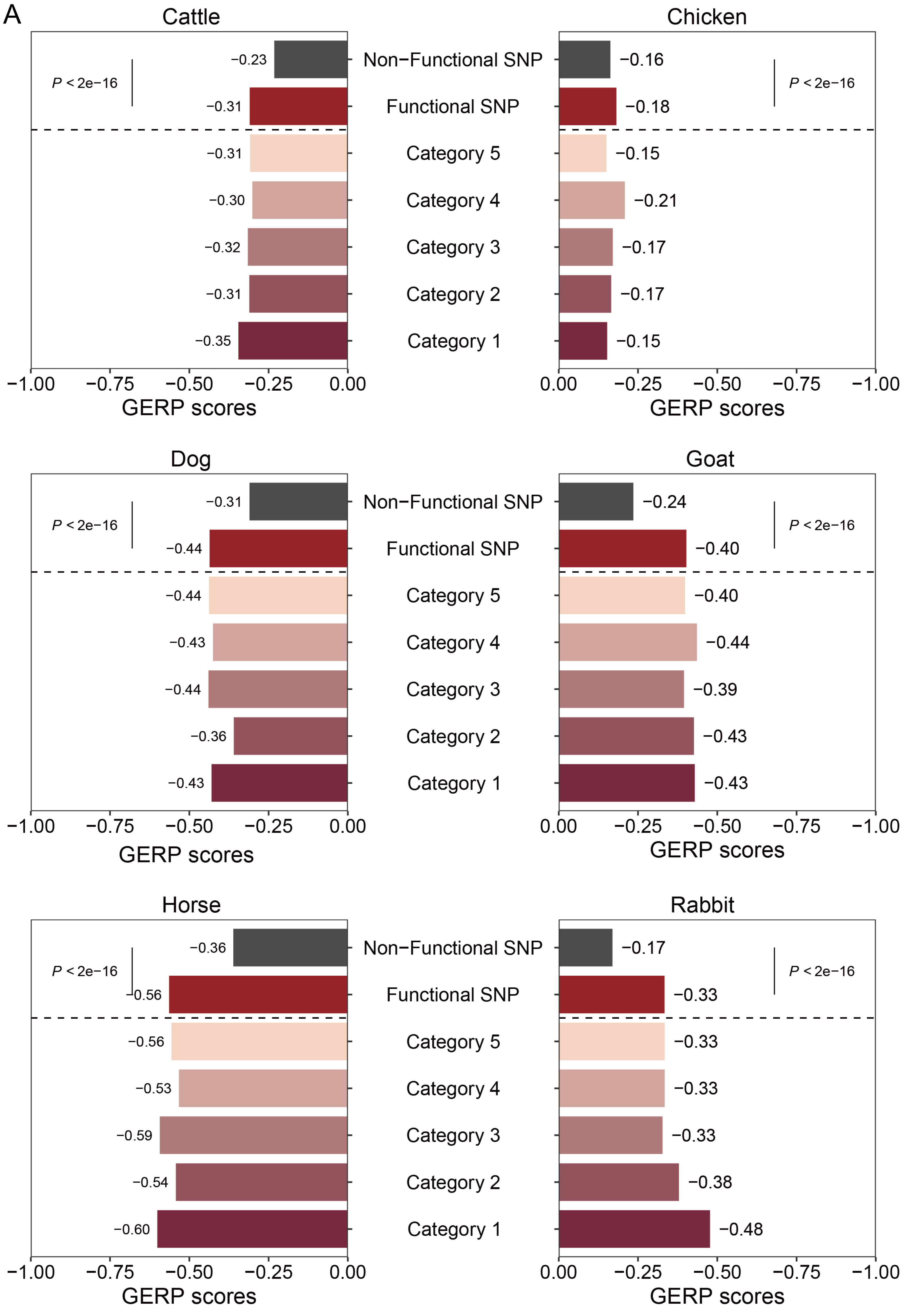

### Supplementary Figure 4

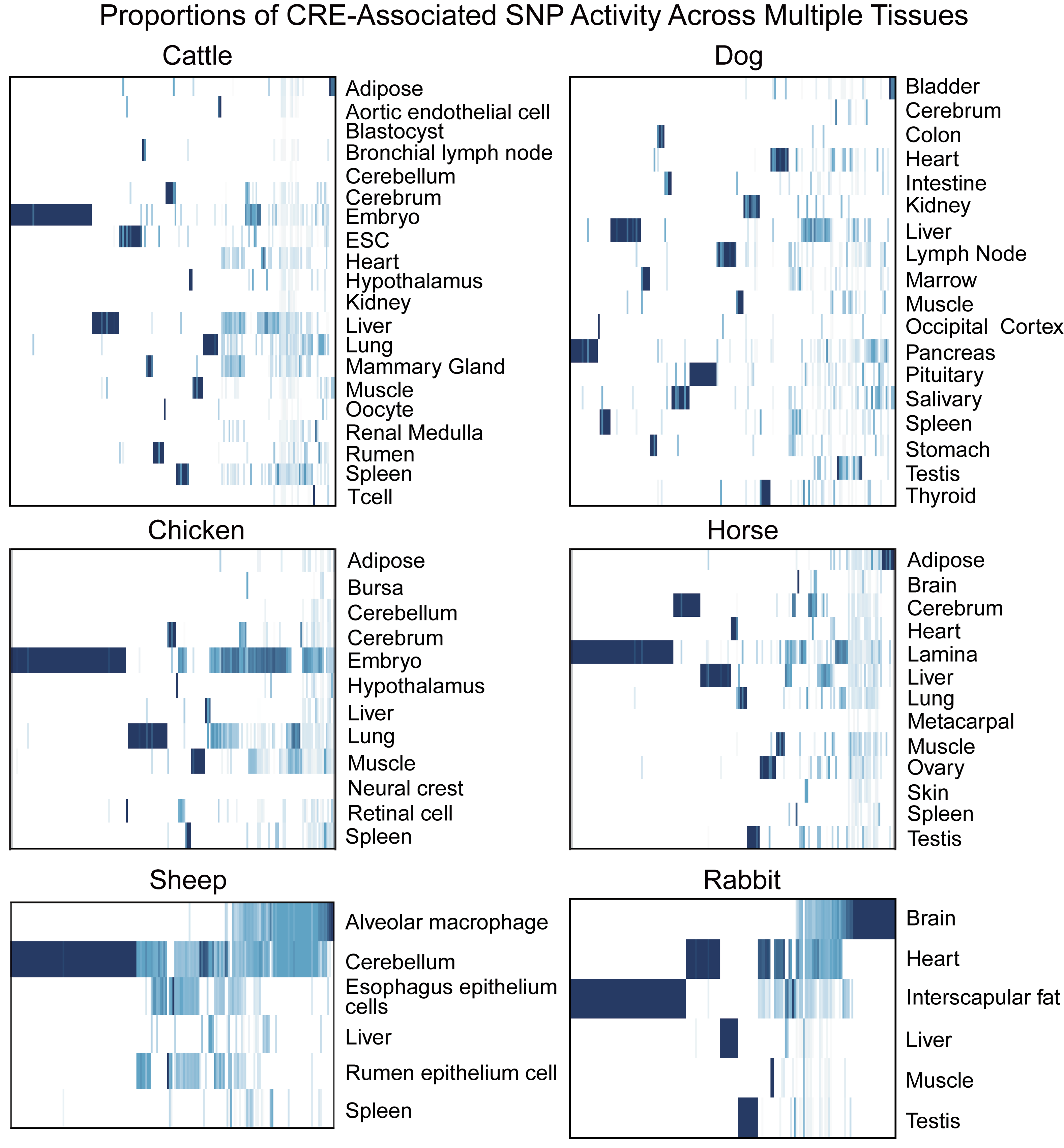
