## Supplementary Figure 5 for "Annotation and assessment of functional variants in regulatory regions using epigenomic data in farm animals"

A

**Category variant by IfmutAnnotator tool**

IFmut — Home | Category | Epigenetics | 3D Genome | Gene | Trait | Single Cell | Visualization | Download | Help

### 3. Category Noval Variants of (Pig)

**Step 1**

Please select file... | Browse | RESET

Users can upload a BED format file containing variant locations to classify novel variants with the IfmutAnnotator tool.
Example: test.bed
Note that the upload file size can not exceed 100kb and line number can not exceed 3000 lines.

**Step 2**

Input

| | | |
|---|---|---|
| chr1 | 7499395 | 7499396 |
| chr4 | 3745800 | 3745801 |
| chr5 | 5964436 | 5964437 |
| chr6 | 5236928 | 5236929 |
| chr9 | 7354473 | 7354474 |
| chr10 | 7347948 | 7347949 |
| chr12 | 7172795 | 7172796 |
| chr14 | 6248770 | 6248771 |
| chr15 | 9027956 | 9027957 |
| chr16 | 7412511 | 7412512 |
| chr17 | 4314012 | 4314013 |
| chr17 | 7258786 | 7258787 |
| chr18 | 4389788 | 4389789 |
| chrX | 6100411 | 6100412 |

**Step 3**

output

| | | | |
|---|---|---|---|
| chr5 | 5964436 | 5964437 | 1b |
| chr4 | 3745800 | 3745801 | 1d |
| chr9 | 7354473 | 7354474 | 2a |
| chr12 | 7172795 | 7172796 | 2b |
| chr1 | 7499395 | 7499396 | 2b |
| chr15 | 9027956 | 9027957 | 3a |
| chr6 | 5236928 | 5236929 | 3b |
| chr17 | 7258786 | 7258787 | 4 |
| chr10 | 7347948 | 7347949 | 5 |
| chr14 | 6248770 | 6248771 | 5 |
| chr16 | 7412511 | 7412512 | Not in category1-5 |
| chr18 | 4389788 | 4389789 | Not in category1-5 |
| chr17 | 4314012 | 4314013 | Not in category1-5 |
| chrX | 6100411 | 6100412 | Not in category1-5 |
